# Alginate-based microparticles coated with HPMCP/AS cellulose-derivatives enable the Ctx(Ile^21^)-Ha antimicrobial peptide application as a feed additive

**DOI:** 10.1101/2021.03.16.435719

**Authors:** Cesar Augusto Roque Borda, Hanyeny Raiely Leite Silva, Edson Crusca Junior, Jéssica Aparecida Serafim, Andréia Bagliotti Meneguin, Marlus Chorilli, Wagner Costa Macedo, Silvio Rainho Teixeira, Elisabete Aparecida Lopes Guastalli, Nilce Maria Soares, Jessica MA Blair, Zoe Pikramenou, Eduardo Festozo Vicente

**Author notes:** **Address for correspondence:** Eduardo Festozo Vicente, São Paulo State University (Unesp), School of Sciences and Engineering, Tupã, São Paulo – Brazil.

## Abstract

Microencapsulation is a potential biotechnological tool, which can overcome AMPs instabilities and reduce toxic side effects. Thus, this study evaluates the antibacterial activities of the Ctx(Ile^21^)-Ha antimicrobial peptide against MDR and non-resistant bacteria, develop and characterize peptide-loaded microparticles coated with HPMCAS and HPMCP. Ctx(Ile^21^)-Ha microencapsulation was performed by ionic gelation with high-efficiency, maintaining the physical-chemical stability. Ctx(Ile^21^)-Ha coated-microparticles were characterized, and their hemolytic activity assay demonstrated that hemolysis was decreased up to 95% compared to single molecule. In addition, *in vitro* release control profile simulating different portions of gastrointestinal tract was performed and showed the microcapsules’ ability to protect the peptide and release it in the intestine, aimed pathogens location, mainly by *Salmonella* sp. Therefore, use of microencapsulated Ctx(Ile^21^)-Ha can be allowed as an antimicrobial controller in monogastric animal production, being a valuable option for molecules with low therapeutic indexes or high hemolytic rates.

## 1. INTRODUCTION

In recent years, consumers have changed perspectives when purchasing a product, focusing mainly on food safety (Heneghan, 2015). This factor is related to the use of nutritionally adequate and safe foods (Coleman-Jensen et al., 2020). When it refers to safe foods, it is basically related to foods that do not affect the consumer health (Chassy, 2010).

Antimicrobial peptides (AMPs) has been a current and highlighted research area because they are not considered drugs and generally do not generate bacterial resistance (Malekkhaiat Häffner & Malmsten, 2019). Generally, AMPs are biologically active with very low concentrations against pathogens such as *Escherichia coli, Staphylococcus aureus* and *Candida albicans* (Lorenzón et al., 2012), being also effective in multidrug-resistant (MDR) bacteria that belong to the ESKAPE pathogen group (Shams Khozani et al., 2019; Yin et al., 2020). These types of peptides were also studied in a large group of Gram-positive and Gram-negative bacteria, exhibiting positive results on *in vitro* (Price et al., 2019) and *in vivo* (Yang et al., 2019) studies.

Ctx(Ile^21^)-Ha, an AMPs extracted from the skin of a Brazilian frog, has emerged as a promising molecule of biotechnological application (Ferreira Cespedes et al., 2012; E. F. Vicente et al., 2013). This peptide demonstrated a high antimicrobial activity against *E. coli, Pseudomonas aeruginosa* (Gram-negative), *S. aureus, Bacillus Subtilis* (Gram-positive) bacteria and fungi *C. albicans, Candida krusei, Candida parapsilosis* and *Cryptococcus neoformans*, which has a great potential of application as a new and natural food additive in animal nutrition. The successful effectiveness of the *in vivo* peptide application depends of the protection or immobilization to avoid nonspecific cellular hemolysis and peptide denaturation/degradation. For this achievement, site-specific carriers such as liposomes, nanoparticles and microparticles can be used (Tagde et al., 2020).

Microencapsulation is a method of protecting bioactive compounds, widely used to preserve and protect their physicochemical and functional properties (Ye et al., 2018). Alginate is a natural polyanionic polymer extracted from seaweed (Lee & Mooney, 2012), used for different encapsulation processes of food products, due to its biodegradability (K. Liu et al., 2006), biocompatibility (S. Wang et al., 2020), dispersive properties (F. Wang et al., 2020) and crosslinking by ion exchange with cationic salts (Dalmoro et al., 2012). Some polymers can be added to microparticles’ development, such as hydroxypropylmethylcellulose acetate succinate (HPMCAS) and hydroxypropylmethylcellulose phthalate (HPMCP). These compounds were largely used as coating, due to their gastric resistance and ease of dissolution in intestine, or at a pH greater than 5.5 (Park et al., 2015). Therefore, they are considered excellent transporters and protectors as coating agents (Momoh et al., 2020), and/or nanoparticle formers (Feng et al., 2019). These aspects motivated the design of a microsystem that leads the antimicrobial peptide to be specifically released in the intestine, to combat common intestinal diseases caused by bacteria such as *Salmonella enteritidis, Salmonella typhimurium* and *E. coli*.

Therefore, if antimicrobial peptides are protected and transported within micropolymeric particles such as alginate and cellulose derivatives (HPMCP/AS), they can be carried to the intestine of any animal of interest (animal production), maintaining their stability and fulfilling their antimicrobial function. In this case, poultry was used as an in vitro monogastric animal model. The goal of this study was to assess the antibacterial activities of the Ctx(Ile^21^)-Ha antimicrobial peptide against MDR and non-resistant bacteria and characterize alginate microparticles containing the antimicrobial peptide Ctx(Ile^21^)-Ha and coated with HPMCAS and HPMCP. The research aimed the protection and improvement/maintenance of physicochemical features of Ctx(Ile^21^)-Ha, also evaluating the microencapsulation hemolytic activities and *in vitro* release profile, for application against gastrointestinal pathogenic bacteria in monogastric animal production, in this way, a new bioproduct was developed with the potential capacity to replace the use of conventional antibiotics.

## 2. MATERIALS AND METHODS

### 2.1. Synthesis, purification and characterization of Ctx(Ile^21^)-Ha antimicrobial peptide

The SPPS detailed methodology of Ctx(Ile^21^)-Ha antimicrobial peptide, also named Ctx, contracted nomenclature used in this work, is described in Vicente et al. (2013) and in the Supplementary Material.

### 2.2. Antibacterial activity assays

The antimicrobial activities of the antimicrobial peptide present new biological activities against *Salmonella enteritidis* (SE), *Salmonella typhimurium* (ST) and multidrug resistant bacteria *Pseudomonas aeruginosa, Acinetobacter baumannii*, and *Staphylococcus aureus*. The essays were carried out in a 96-well ELISA tray, following the microdilution method (Wikler, 2006). *Salmonella* sp. were isolated from laying hens with microbiological tests for strains’ confirmation. For clinical resistant bacteria, growth colonies were made from original bacteria strains.

### 2.3. Production of coted-microparticles (ER)

The microparticles were obtained using ionic gelation of sodium alginate in aluminum chloride, as previously described (Deladino et al., 2008) and coated with HPMCP (H) and HPMCAS (C) (Supplementary Material).

### 2.4. Characterization of coated-microparticles

For the characterization of the microparticles, tests were performed: Encapsulation efficiency, FT-IR/FT-Raman, Morphological, Thermal and X-ray diffraction analysis (Supplementary Material).

### 2.5. *In vitro* release profile

*In vitro* release study analysis were performed in two stages, according to the Morgan et al. (2020), with some modifications: i) *simulated gastric fluid (SGF)*, procedure that mimics the proventriculus and gizzard (considered chemical and mechanical poultry stomachs). A mass of 50.8 ± 3.54 mg of microcapsules were placed in a SG solution at 42°C, for 20 min (Supplementary Material).

### 2.6. Hemolytic activity assay

The hemolytic activity assay of pure and microencapsulated peptides was determined by simulating the digestion time, based on the analysis of hemoglobin release from erythrocytes (Serrano et al., 2013) (Supplementary Material).

## 3. RESULTS AND DISCUSSIONS

### 3.1. Ctx(Ile^21^)-Ha antimicrobial peptide analysis

The detailed synthesis, chromatogram profile and mass spectra are presented (Fig. 1) in the Supplementary Material.

**Fig. 1.**
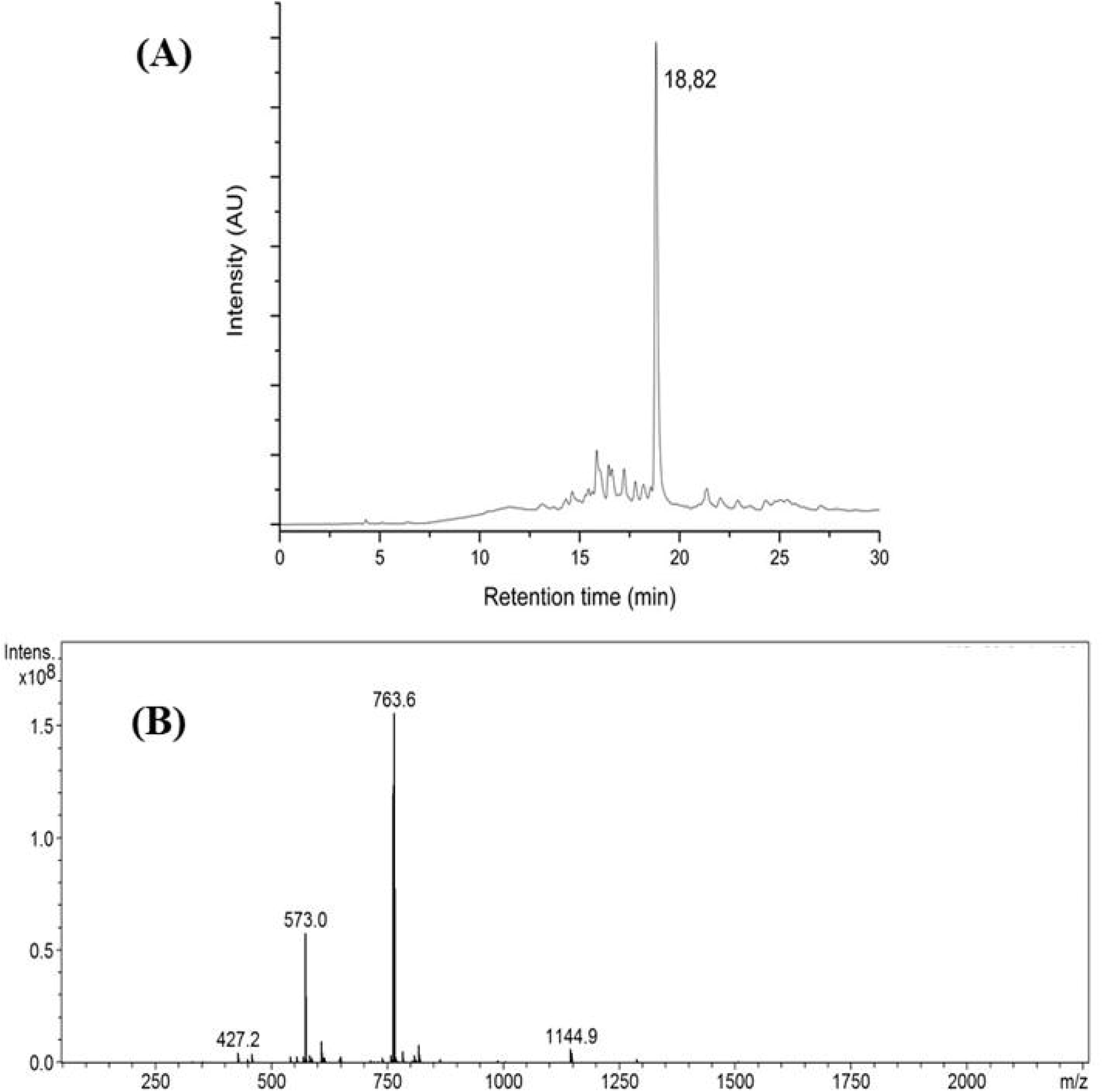
(A). HPLC analysis of crude antimicrobial peptide Ctx(Ile^21^)-Ha. **(B)**. Mass spectra of the antimicrobial peptide Ctx(Ile^21^)-Ha, confirming the correct characterization.

### 3.2. Antimicrobial Activity

The results of the antimicrobial activity assay are showed in the Table 1. It is possible to note that the Ctx(Ile^21^)-Ha peptide was active against both strains of *Salmonella* pathogens tested. The MIC is notably low (4 µmol L^-1^) against the MDR and Gram-positive bacteria *Staphylococcus aureus*, also exhibiting antimicrobial activity with the others Gram-negative bacteria tested, including the MDR *P. aeruginosa* and *A. baumannii*. In addition, the molecule showed a remarkably antimicrobial activity against MDR bacteria, whose results were better than the commercial antibiotic (ampicillin) used as positive control. According to these results, this assay could allow the microencapsulation development of peptide Ctx(Ile^21^)-Ha, since it showed very active against the most hazardous multidrug resistant bacteria.

Other AMPs were studied to inhibit the replication of *Salmonella* sp., such as cathelicidin-BF, Cathelicidin-BF, which demonstrated an anti-ST effect in infected murines, however its application was as injectable, a process that is to be avoided in animal production, and no values for anti-SE were shown, however, the challenges in the transport of the peptide made this treatment not entirely efficient (Xia et al., 2015). Ctx microparticles being the ones that showed the greatest advantage as they were a product for oral administration. Ctx also has anti-SE MIC values of 4 µmol L^-1^, better than indolicidin (8.4 µmol L^-1^), so its potential is maintained. Furthermore, human cathelicidin LL-37, a highly studied peptide with potential activity, showed low anti-ST activity (28 µmol L^-1^), but did not show anti-SE activity. Ctx showed better anti-SE results compared to conventional drugs such as Chlortetracycline (36 µmol L^-1^) and Neomycin (13 µmol L^-1^) (Y. Liu et al., 2011). Which shows the efficiency of this peptide against the problem posed and due to the lack of published articles on HPMCP/AS-coated microparticles using ionic gelation with antimicrobial peptides, we proposed a novel product with combined techniques from the food and pharmacy industries.

### 3.3. Microencapsulation analysis

EE was considerably high, reaching values higher than 70%, as shown in Table 2 (standard deviation margin for this data set was less than 0.001). This yield is related to the polymer concentration and interaction between peptide and polymer (Yeo & Park, 2004). In case of proteins and peptides, the efficiency would also be related to amphipathic effect that the Ctx peptide has, since the hydrophobic interaction is a dominant force between peptide and polymer. In this manner, these compounds can perform an ion exchange that optimally encapsulates the polymeric network mesh that contains −COOH groups (Jyothi et al., 2010). Based on these antecedents, it is evidenced that the power of incorporation of the encapsulating matrix with the Ctx peptide was adequate for an effective encapsulation.

As expected, microcapsules obtained by small droplets incorporated a small fraction of the peptide, so the concentration also decreased. However, the principal component analysis (PCA) indicates that their efficiency is maintained, except for C2 (P<0.001). These findings indicate that the coating procedure was carried out correctly and microcapsules show the necessary solidity for do not break or dissolve instantly. In addition, these systems will only release the Ctx peptide after HPMCAS and HPMCP degradation (Momoh et al., 2020).

#### 3.3.1. Fourier transform spectroscopy analysis

The bands and peaks of the polymers in question were considered, comparing them with sodium alginate, HPMCP, HPMCAS and the synthesized antimicrobial peptide isolated. In Fig. 2A and 2B, the characteristic peaks of aromatic amino acids, such as tryptophan and phenylalanine at 1,136, 1,180 and 1,202 cm^-1^ are observed (Ayodhya et al., 2015; Ciobanu et al., 2015), and peaks of ∼1,650 cm^-1^ (C=O elongation) and ∼1,540 cm^-1^ (N-H bending vibrations and C-N elongation of the main peptide axis) which indicate the presence of amide I and amide II bands, respectively, typical of all amino acids present. In addition, the band observed at 1,620 cm^-1^ can be related to the hydrophobic nature of the peptide (Miller et al., 2013). The band at ∼1,457 cm^-1^ can be related to –CH_3_, –CH_2_ from aromatic rings and/or β-sheet of the peptide secondary structure. These bands are related to aromatic amino acids, flexural vibration (δ) of C–H product of the alkyl chains and aromatic stretching vibration (Miller et al., 2013; Oliveira et al., 2016).

**Fig. 2.**
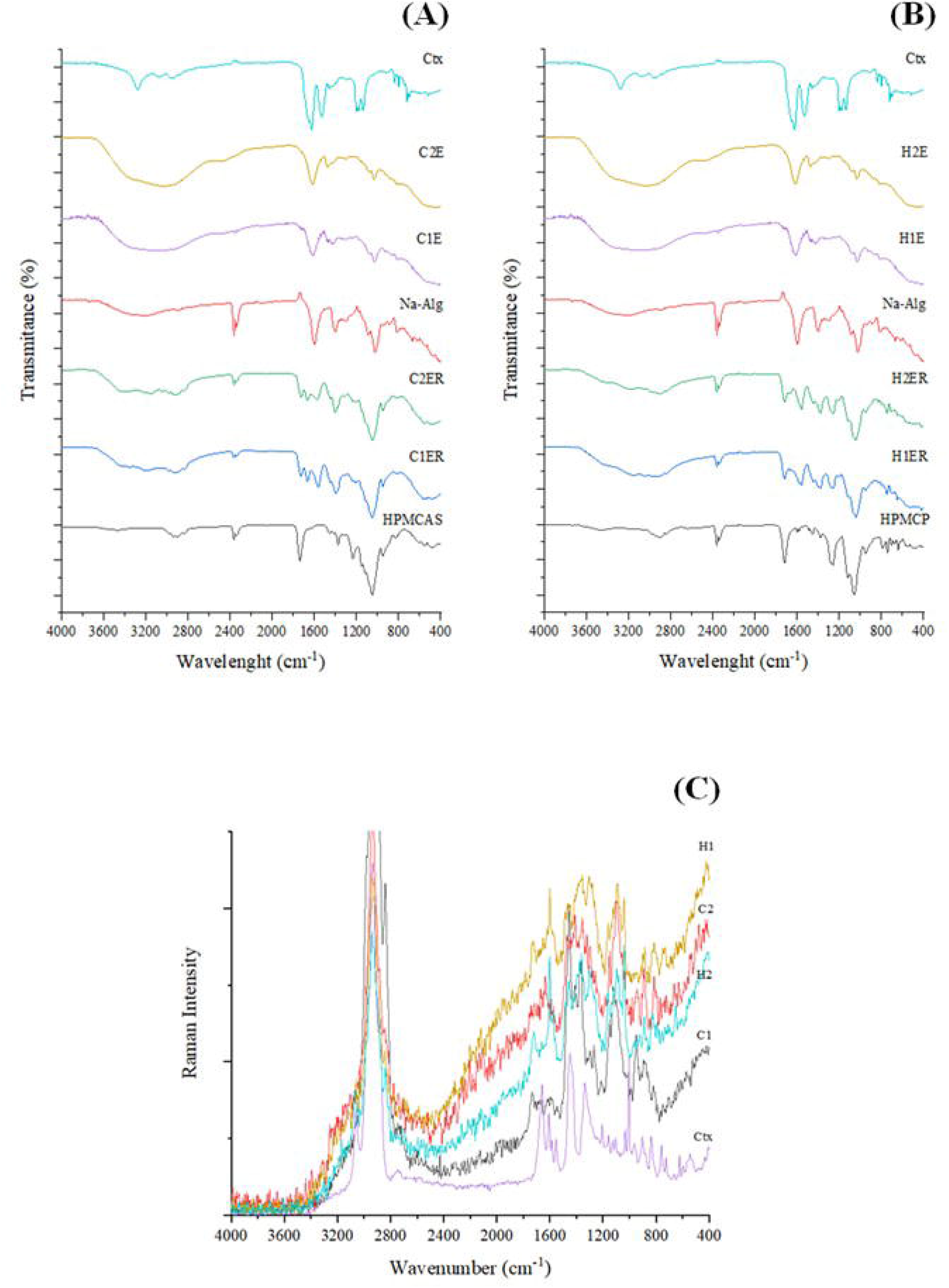
Ctx(Ile^21^)-Ha peptide and microencapsulation phases. **(A)** FTIR-ATR spectra of HPMCAS. **(B)** FTIR-ATR of HPMCP. **(C)** FT-Raman spectra of HPMCAS and HPMCP.

The FT-Raman spectrum confirmed the presence of the Ctx(Ile^21^)-Ha peptide (Fig. 2C) by the existence of aromatic amino acids bands, such as tryptophan, at 760 cm^-1^ (weak intensity), 885 cm^-1^ (weak), 1,616 cm^-1^ (medium); phenylalanine at 622 cm^-1^ (weak), 1,333 cm^-1^ (strong), 1,585 cm^-1^ (weak), 1,605 cm^-1^ (medium), and, finally, a band at 1,004 cm^-1^ (strong) representing both amino acids. Other results confirm the mentioned observations as demonstrated in Shao et al. (2005).

The peptide secondary structure characteristics were also found: i) Amide I: C=O and C=C elongation and the α-helix secondary structure detected at 1,661.5 cm^-1^ (very strong); ii) Amide III: strain (CH_2_), 1,299.5 cm ^-1^ (shoulder); β sheet at 1,232 cm^-1^ (weak) and β and disordered sheet at 1,264 cm^-1^ (weak). Also, in 1,447 cm^-1^ (very strong), a bending mode of CH_2_ and CH_3_ was found (Balgoon et al., 2018). In this way, the FT-Raman technique helped to verify if there was any interference with the water in the FTIR analyzes, since the water does not interfere with the light scattering of the FT-Raman (E. Vicente et al., 2015).

### 3.4. Morphological analysis

In the morphological analysis, it was possible to visualize the microcapsule structures, confirming the formation and thickness of the coating (Fig. 3). HPMCP coating (H1 and H2) showed thickness of 35.8 ± 8.2 µm and 46.7 ± 12.2 µm, respectively. In parallel, HPMCAS coating (C1 and C2) exhibited thickness of 44.6 ± 13.5 µm and 48.4 ± 1.1 µm, respectively. The photomicrographs demonstrated a homogeneous peptide entrapment within the microcapsule structure. Also, it can clearly observe the difference between the blank capsules that contain only sodium alginate, revealing uniform and completely homogeneous branches, in according with some authors (de Farias et al., 2018; Hartmann et al., 2010). However, the surface-formed microcapsules do not show a circular shape as found on their initial shape (data not shown), effect caused by water decrease after drying.

**Fig. 3.**
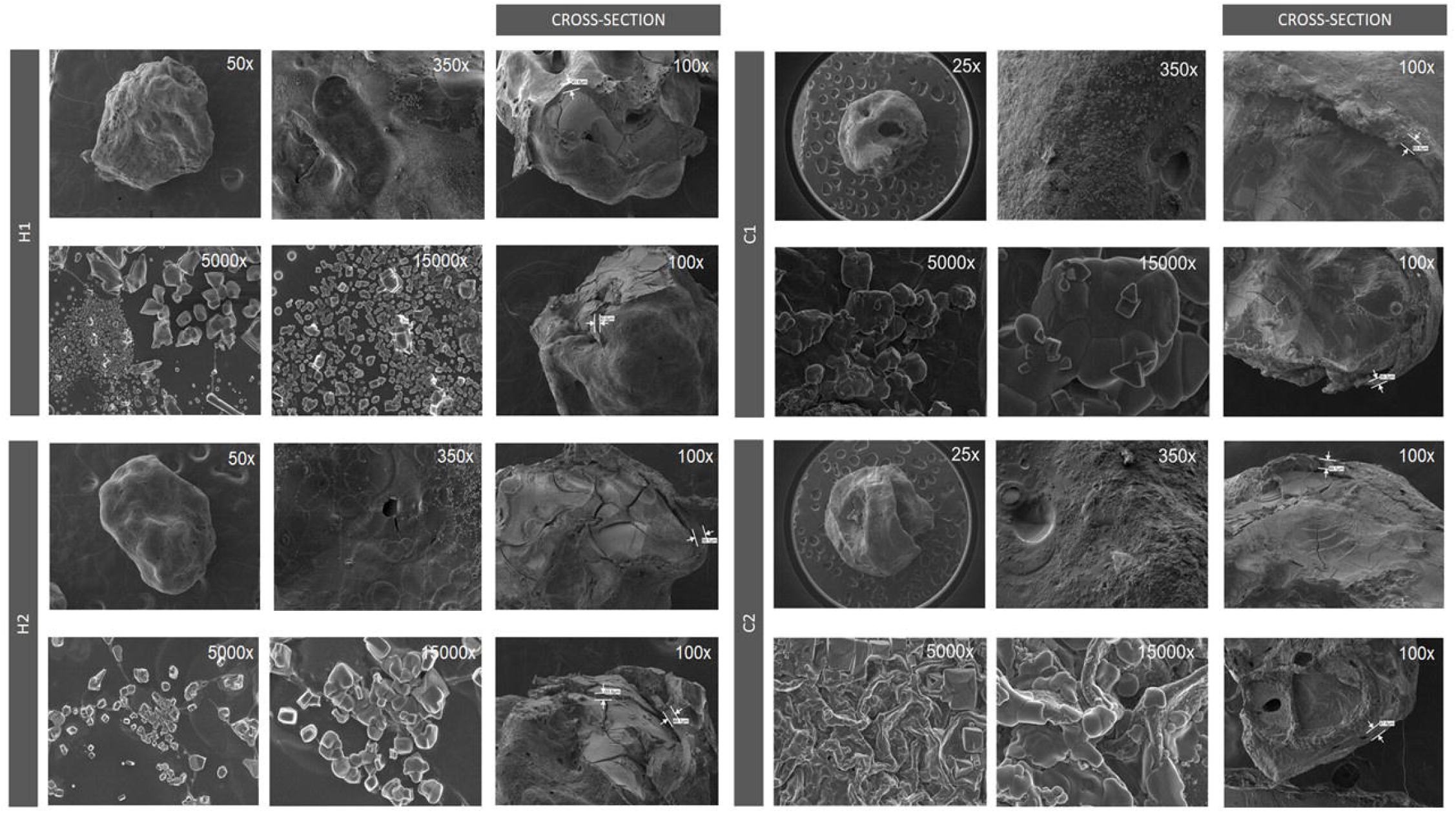
Photomicrographs of the coated microparticles loaded with Ctx(Ile^21^)-Ha peptide. The optical magnitudes vary from 50, 350, 5000 and 15000 x. The cross section makes it possible to confirm the presence and thickness of the coating for each designed microsystem.

Other studies indicated that encapsulated formulations made with sodium alginate presented deformations after drying and a microstructural perfection is not achieved. However, controlled release efficiency of the active principle was carried out normally (Lupo et al., 2015). The use of pharmaceutical combinations such as sodium alginate/soy protein provided rigidity, obtaining spherical capsules. Also, it was evidenced that this effect would be related to the concentration ratio of each encapsulating agent (Volić et al., 2018), as well as the concentration of the crosslinking solution, which in this case was aluminum chloride (Lupo et al., 2015). Moreover, it was noted that some photomicrographs showed the presence of crystals on the surface. This can be related to the presence or formation of sodium chloride salts, attributed to the ionic exchange produced in ionic gelation (Deladino et al., 2008).

### 3.5. Thermal results

The loss of mass due to an increase in temperature is a problem for many peptides and proteins studied. It is caused by the denaturation that it undergoes in the structure, it is generally an irreversible process. Thermogravimetric analysis (Fig. 4) helped to analyze the loss of material, expressed as a percentage of mass. Interestingly, the opposite occurs if the glass transition (Tg) is found, whose denaturation process is reversible (Ricci et al., 2018). However, the Ctx peptide shows no weight loss below 100°C. In addition, samples present a similar behavior, so there is no weight loss in the range that represents food products. However, the encapsulated ones present Tg in the same range, which is a natural effect of the polymers used as reference (sodium alginate, HPMCP and HPMCAS).

**Fig. 4.**
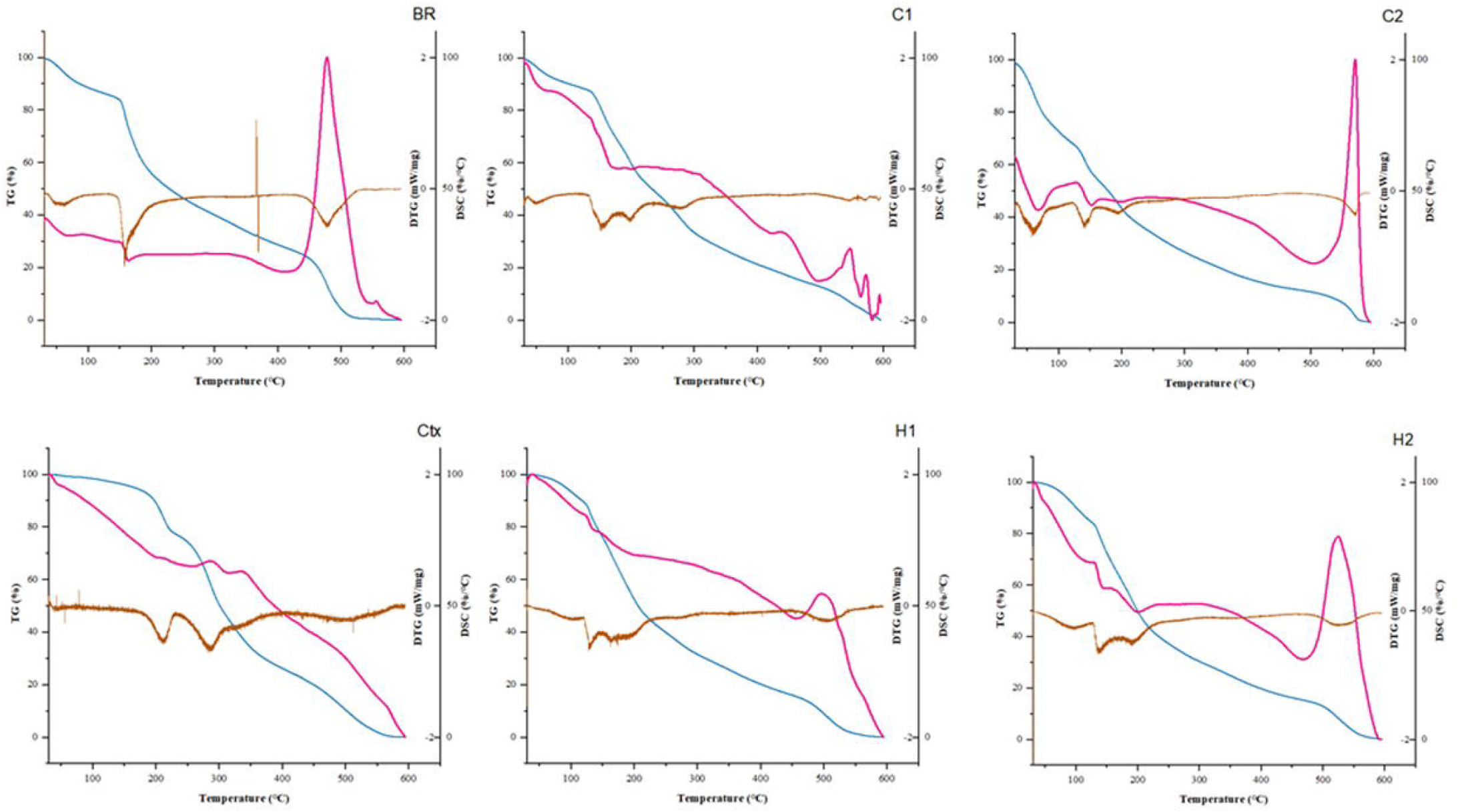
Analysis of TGA (blue), DSC (pink) and dTGA (brown) of all samples.

As shown in the Fig. 4, the first derivative of TGA (dTGA) identifies the thermo-oxidation reactions caused by increments in temperature. Thus, the pure Ctx peptide showed two events between 200 and 300°C (event that, for food products, is not relevant). However, by joining the peptide with the polymers through microencapsulation, an adaptive behavior is observed, showing thermal degradation events of alginate + HPMCAS (Fig. 4BR). Hence, it is suggested that HPMCAS fails to protect the thermal degradation effect of alginate as HPMCP does (Fig. 4H1 and 4H2). In addition, the capacity of energy retention of the polymers used in microencapsulation could significantly destabilize the peptide and some polymers (Tang et al., 2006).

In the case of C2, the difference in the behavior of thermal degradation is remarkable, showing weight decrease below 100°C. It is presumed that the exposure of the dispersion solution for a longer time to achieve complete homogenization, weakening the alginate bonds, which, when undergoing an excess of temperature, cannot remain fixed. Therefore, it is necessary to be careful with the stirring time when preparing microencapsulates.

The glass transition temperature (Tg) is an important parameter to evaluate the microcapsules status during their storage, which makes it possible to visualize a significant difference between the BR and C2 capsules compared to the others products (Lupo et al., 2015). This would indicate that there is a greater stability to the C1, H1, and H2 capsules at room temperature and, as the temperature increases, particles interactions and diffusion also accelerates (Deladino et al., 2008).

In summary, the microencapsulated-coated the antimicrobial peptide by methods applied in this work are semi-tolerant at high temperatures, but thermostable at temperatures that are crucial in the food industry.

### 3.6. XDR analysis

According to the diffractograms (Fig. 5), the antimicrobial peptide Ctx(Ile^21^)-Ha has intensities of 2θ with characteristics of proteins and peptides, located in the 10° band, which represents the α-helix secondary structure. In addition, in the 20° band, which is related to the structure of the β sheet, showing that the peptide has a semi-amorphous solid structure. This effect is related to the peptide solubility, which contains randomly disposed and easily separated molecules (Mendes et al., 2012; Zhao et al., 2015). The use of polymers as bioactive protectants is commonly employed in the pharmaceutical industry, since it helps to reduce drug crystallinity and, in turn, increases their stability (Al-Obaidi & Buckton, 2009), effect that can also be observed in the previous SEM micrographs, confirmed by FT-IR and FT-Raman data. The microcapsules showed typical peaks of alginate and cellulose (HPMCP and HPMCAS conformation). In addition, microencapsulation has a direct relationship with the decrease in the intensity of the crystalline region of the peptide, which facilitates its dissolution and controlled release (Mendes et al., 2012).

**Fig. 5.**
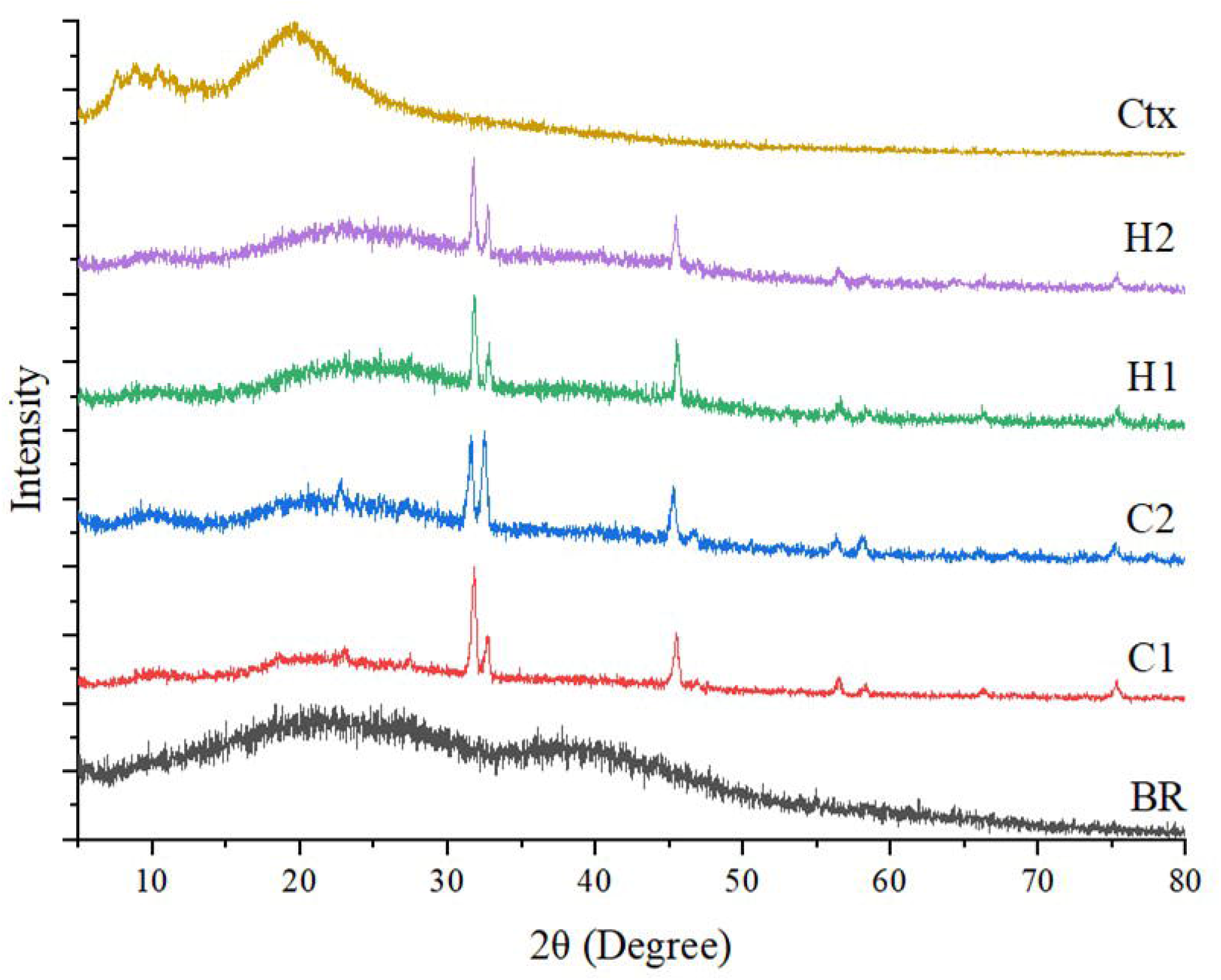
XRD of the samples synthesized and microencapsulated using ionic gelation and fluidized-bed combined method.

### 3.7. *In vitro* release profile

The release profile of peptide from microcapsules was built in two stages, following the gastric and intestinal portions of GIT. First, about 7.74 ± 0.11% and 14.9 ± 0.03% of peptide was released at SGF from H1 and H2 (HPMCP coated microcapsules), respectively. These values indicate that coating with HPMCP makes the microcapsules resistant in an acid environment as proposed (Fig. 6). The higher release rates from H2 compared to H1 are due to the greater amount of peptide encapsulated in these samples (0.4 g L^-1^ Ctx *versus* 0.2 g L^-1^ Ctx), contributing to the creation of a greater concentration gradient that favors diffusion.

**Fig. 6.**
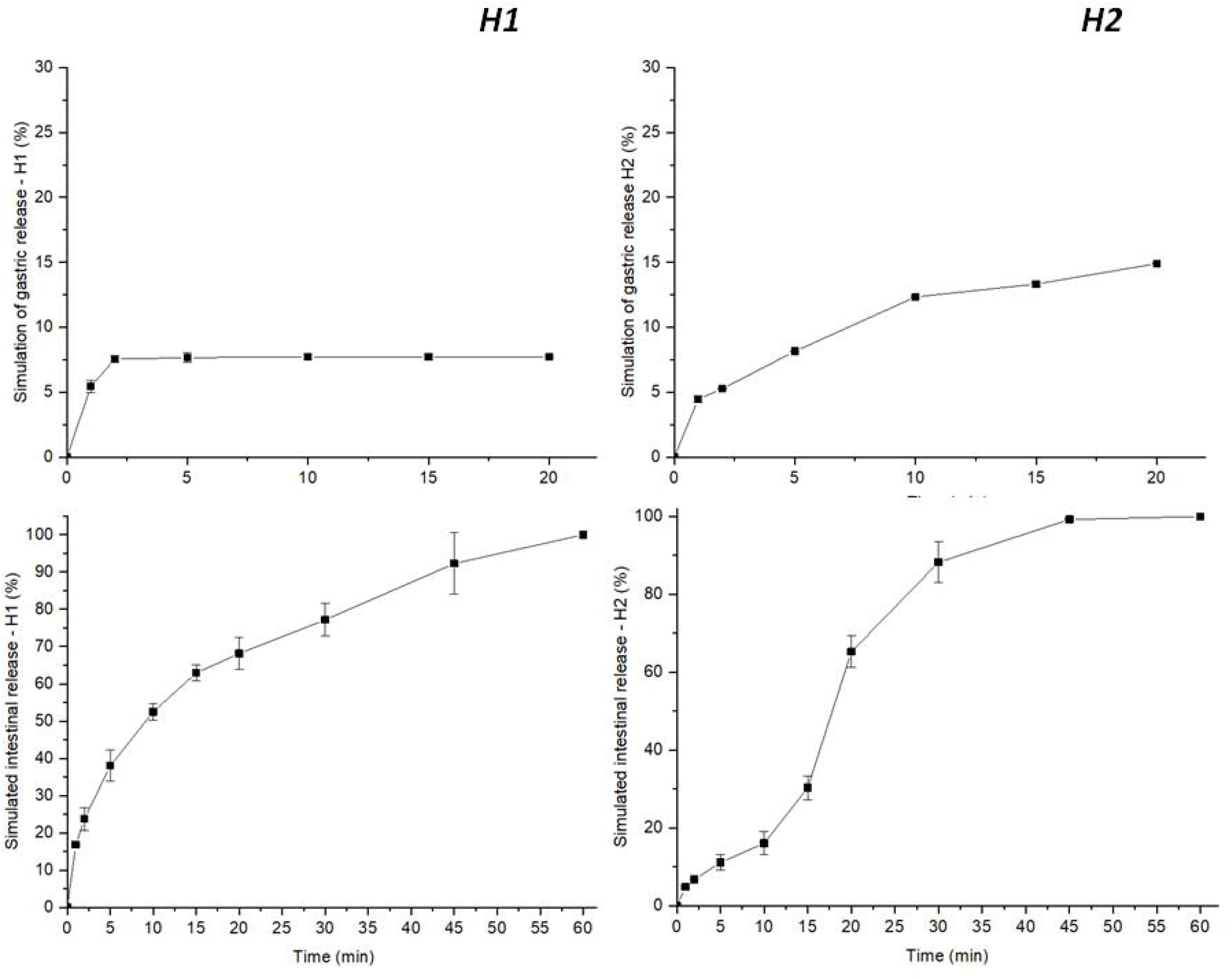
Controlled release of simulated gastric and intestinal systems in laying hens for Ctx encapsulated and coated with HPMCP (data are means ± s.d., n = 3).

On the other hand, microcapsules coated with HPMCAS showed a peptide release of up to 33.31 ± 1.35% for C1 (Fig. 7). As expected, C2 released more than half of the peptide content in a short period, which can be explained by an incomplete a defective microcapsule coating.

**Fig. 7.**
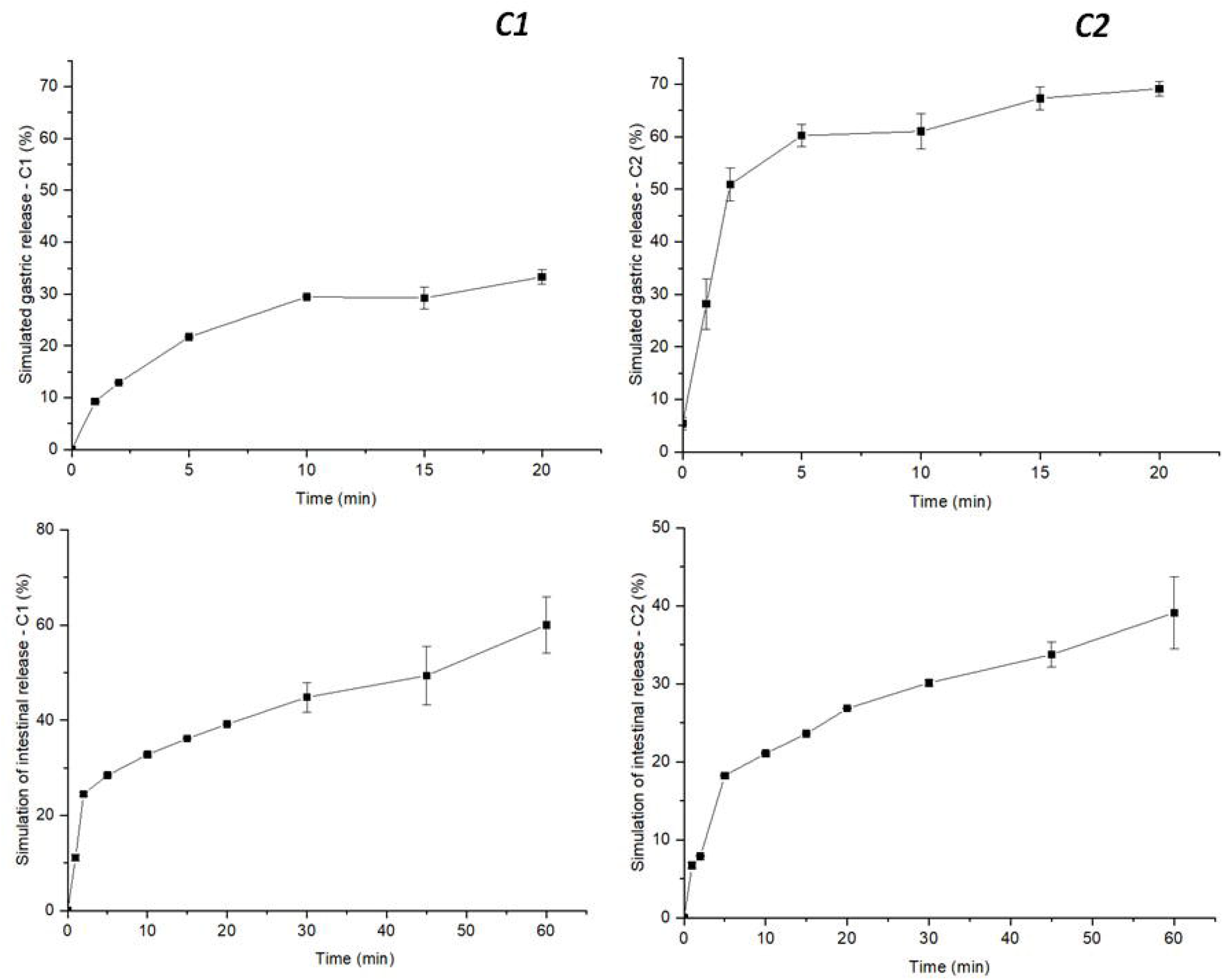
Controlled release of simulated gastric and intestinal systems in chickens for Ctx encapsulated and coated with HPMCAS (data are means ± s.d., n = 3).

Some authors described encapsulated products with similar protective effects (controlled release with HPMCP and HPMCAS) in the food industry (Ahn et al., 2019; Son et al., 2016). In Animal Science, a study was developed producing spheres loaded with Porcine Epidemic Diarrhea Virus (PEDV) to induce immunity by generating antibodies to combat this virus. These spheres were coated with HPMCP to achieve their release, which exhibited positive and similar to our results, indicating that PEDV was slightly released in the gastric system, and entirely in the intestine (Wen et al., 2018). Another study based on the microencapsulation of lactase for dairy products, was successfully carried out releasing around 0.2% and 90% in simulated gastric and intestinal fluid (SGF and SIF, respectively) of the enzyme (Ahn et al., 2019). On the other hand, a study with HPMCAS-coated and encapsulated insulin tended to release around 20% in SGF results, which resemble C1 capsules data, and up to 90% in SIF, similar to all others results presented in this study (Momoh et al., 2020). It has also been reported that formulations with HPMCAS inhibit the presence of enzymes such as trypsin, a peptidase produced in the pancreas and secreted in the duodenum (part of the intestine) (Mcmanus et al., 2020). Likewise, drugs coated with cellulose-derived polymers would provide greater bioavailability and better pharmacological action of the coated products (Maderuelo et al., 2019). In summary, HPMCP and HPMCAS coated products tend to slightly release the bioactive compound in the digestive tract and release completely in the intestinal tract, maintaining its biological activity and its physicochemical stability.

The effect of peptide intestinal release for all microcapsules was efficient when exposed in the intestine fluid environment, which it is the preferred site of bacterial multiplication, mainly of *Salmonella enteritidis*. Thus, the release was carried out satisfactorily in a non-acid medium, which can allow the Ctx peptide to be active against the pathogenic bacteria present in this organ. HPMCP is a polymer responsible for protecting gastro-sensitive molecules, since it is insoluble at pH ∼ 3 and, consequently, also protects from the enzymatic and oxidative processes in this environment (Chung et al., 2014). This compound is a modification of cellulose, where the carboxyl groups were replaced by hydroxyl groups, which gives it a high solubility at pH ∼ 7. This feature allows coated molecules to have stability during transport at a lower pH (Surini & Prakoso, 2018). On the other hand, HPMCAS is also a biodegradable polymer with similar properties. However, the ratio of acetyl and succinyl groups would be the main source of solubility of this polymer, the pH being able to range between 5.5 to 6.5 (Fu et al., 2020).

These plasticizing polymers normally generate cracks as a dry final product after coating by fluidized bed. For this reason, coating preparation was improved with triethyl citrate, which gave greater firmness and remarkedly decreased cracks, effect that is corroborated with photomicrographs of the SEM (Deshpande et al., 2018). The influence of compounds that improve enteric coating based on compounds derived from hydroxypropyl methylcellulose (HPMC) is not yet clear. Although, it was recently reported that the combination of talc/triethyl citrate as an additive could be added to close these cracks (Fu et al., 2020). However, these drawbacks do not interfere with the release of the peptide in the intestinal tract positively, which is the main goal of the Ctx(Ile^21^)-Ha peptide microencapsulation.

### 3.8. Hemolytic activity

According to previous published studies (E. F. Vicente et al., 2013), the hemolytic activity (HC_50_) of the antimicrobial peptide Ctx(Ile^21^)-Ha is 7.1 µmol L^-1^. The results obtained from this study indicate that the level of hemolysis of the peptide was controlled and strongly reduced, with a rate greater than 95% with the microencapsulation process. Thus, the present analysis showed that, in the simulated period of digestion, according to Morgan and collaborators (Morgan et al., 2020), hemolytic activity of the microparticles were below 10%. Therefore, the present coated-microcapsules do not have hemolytic activity, which represents a positive feature regarding the peptide toxicity. The kinetics of hemolytic activity are shown in Fig. 8.

**Fig. 8.**
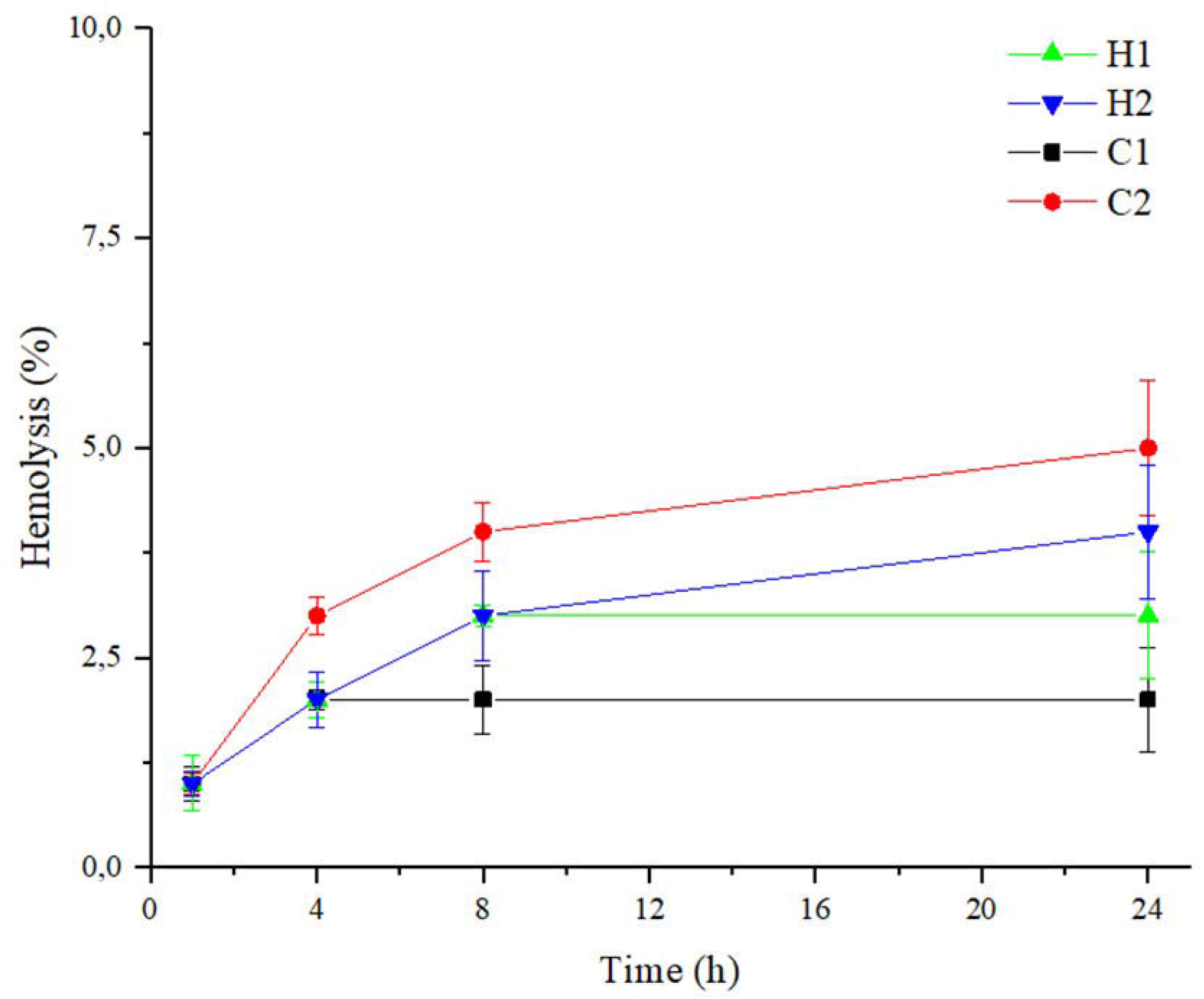
Evaluation of the hemolytic activity of all samples to determine the viability of the product and its application as a feed additive.

## 4. CONCLUSIONS

The antimicrobial peptide Ctx(Ile^21^)-Ha shows a broad and powerful antimicrobial activities against most hazardous pathogens in poultry production and multidrug resistant bacteria, which can be applied as an efficient natural antibiotic. This work demonstrated that Ctx(Ile^21^)-Ha was properly microencapsulated and coated with HPMCAS and HPMCP, confirming its characterization using several physicochemical techniques. In addition, peptide physicochemical properties in the microcapsules were maintained stable and, in some case, optimized, as hemolytic activity decreased by more than 95%, which became the formulation secured and stable. *In vitro* analysis of Ctx microparticles showed that controlled release helps to maintain and protect the peptide against different events that could alter its structure, offering the release of the entire peptide in an intestinal pH. From the best of our knowledge, the present work is a pilot study for development of a new and efficient animal feed additive, since there are no studies of microencapsulated antimicrobial peptides coated with HPMCP or HPCMAS coating have been reported yet. Therefore, the use of microencapsulated antimicrobial peptide Ctx(Ile^21^)-Ha can be an interesting option as a possible natural growth promoter, opening opportunities to replace the using of synthetic antibiotic additives in animal feeding.

## Declarations of competing interest

All contributing authors declare no conflicts of interest.

## CRediT authorship contribution statement

**Cesar Augusto Roque Borda:** Writing -original draft, Investigation, results analysis and performing of all the tests and applied techniques. **Hanyeny Raiely Leite Silva:** peptide synthesis and microencapsulation. **Edson Crusca Junior:** hemolytic activity assays, English corrections and article review. **Jéssica Aparecida Serafim:** analytical profile and HPLC. **Andréia Bagliotti Meneguin and Marlus Chorilli:** encapsulation and coating process design. **Wagner Costa Macedo and Silvio Rainho Teixeira:** microparticles characterization by DSC/TGA and XRD. **Elisabete Aparecida Lopes Guastalli, Nilce Maria Soares, Jessica MA Blair and Zoe Pikramenou:** antibacterial analysis and article review. **Eduardo Festozo Vicente:** Conceptualization, Development, Supervision, Funding acquisition, article review and writing.

## Acknowledgments

This work is supported by São Paulo Research Foundation/FAPESP (Process number 2016/00446-7) and master scholarships (Proc. 2018/25707-3 and 2018/25735-7). SEM FEG microscopy laboratory at Institute of Chemistry – Araraquara (UNESP) for the micrographs; University of Araraquara which made available the use of the fluidized-bed equipment and Shin-Etsu company for gently donate and provide the HPMCP and HPMCAS coatings for the experiments. Finally, the research group “Peptides: Synthesis, Optimization and Applied Studies -PeSEAp”. This work is a chapter of master’s thesis and is also part of the national patent protected throughout the Brazilian territory by the National Institute of Intellectual Property (INPI BR1020200220489).

## Notes

### Competing Interest Statement

The authors have declared no competing interest.

