## SUPPLEMENTARY MATERIAL for "Alginate-based microparticles coated with HPMCP/AS cellulose-derivatives enable the Ctx(Ile^21^)-Ha antimicrobial peptide application as a feed additive"

**MATERIAL AND METHODS**

2.1. **Chemical reagents**

Hypromellose acetate succinate (HPMCAS, AQOAT® - Grade AS-LF) and hypromellose phthalate (HPMCP, Grade HP-55, nominal phthalyl content 31%) coatings were kindly donated by Shin-Etsu Chemical Co.,Ltd. – Japan, sodium alginate low molecular weight (12,000–40,000 g mol^−1^, M/G ratio of 0.8) and the other chemical reagents were purchased at HPLC grade from Sigma-Aldrich Co. (MO, USA). Fmoc-amino acids, resins (Rink amide MBHA), trifluoroacetic acid (TFA), 4-methylpiperidine, 1-hydroxybenzotriazole hydrate (HOBt), triisopropylsilane (TIS), *N,N*′-diisopropylcarbodiimide (DIC) and acetonitrile were obtained in HPLC/analytical grade from AAPPTEC (Louisville, KY, USA), dimethylformamide (DMF) was purchased from Neon Comercial (São Paulo, Brazil), and dichloromethane (DCM) was purchased from Anidrol Products Laboratories (São Paulo, Brazil).

2.2. **Synthesis, purification and characterization of Ctx(Ile^21^)-Ha antimicrobial peptide**

**2.2.1.	Synthesis of Ctx(Ile^21^)-Ha antimicrobial peptide**

The 21-mer antimicrobial peptide Ctx(Ile^21^)-Ha (Gly-Trp-Leu-Asp-Val-Ala-Lys-Lys-Ile-Gly-Lys-Ala-Ala-Phe-Asn-Val-Ala-Lys-Asn-Phe-Ile), also named Ctx, contracted nomenclature used in this work, was assembled by standard solid phase peptide synthesis methodologies assisted manually (Fmoc/tBu) (E. F. Vicente et al., 2013). Briefly, the solid support (Rink amide MBHA resin) was previously washed with 1:1 v/v DMF/DCM. A solution of 20% 4-methylpiperidine in DMF was used to remove the Fmoc group. The next Fmoc-amino acid (1.2 eq, molar equivalents) was coupled with 1.2/1.2 eq of HOBt/DIC in DMF/DCM for 2 h at room temperature (RT). The Kaiser test (Kaiser et al., 1970) was used to monitor the couplings, analyzing directly the free amino groups. This fast but destructive essay indicates whether the peptide is protected (yellow color - absence of free amino groups) or unprotected from the Fmoc protector (blue color - presence of free amino groups) and also to confirm if the coupling was successful. For the test, some very few resin beads were placed with two drops of ethanolic solutions (5% v/v ninhydrin and 80% w/v phenol) and pyridine solution (2% v/v of 10^-3^ mol L^-1^ of potassium cyanide) at 120°C for 4 min.

After complete coupling of peptide primary sequence, the peptide was removed from the resin using a solution composed of TFA, TIS and water (95:2.5:2.5, v/v/v) for 2 h at room temperature (RT). Subsequently, peptide was precipitated with cold ethyl ether and centrifuged (5 min at 6,500 rpm at RT, in triplicate), to separate by-products from resin cleavage reaction, which are miscible in this solvent. Then, ether supernatant containing the hydrophobic by-products was reserved and the solid peptide/resin phase was dried to remove all remaining ether by evaporation. In dry samples, a solution containing aqueous solvent (A, 0.045% TFA) and acetonitrile solvent (B, 0.036% TFA, 1:1, v/v) was added and centrifuged, obtaining the resin (solid phase) and the peptide (liquid phase). The samples were lyophilized for 48 h in a lyophilizer (Liotop, model K108, Brazil), and their purity was evaluated by liquid chromatography (next section).

**2.2.2.	Purification and characterization of Ctx(Ile^21^)-Ha peptide**

The purity degree and confirmation of the antimicrobial peptide Ctx(Ile^21^)-Ha were determined by HPLC (Shimadzu, with DGU-20A5R membrane degasser, CTO-20A column oven, sampler automatic SIL-10AF, fraction collector FRC-10A, UV detector SPD-20A and LC-20AT of double pump) and Mass Spectroscopy. Initially, an analytical analysis was performed to obtain the crude peptide profile, the system was equipped with a C18 reverse phase column (dimensions of 250 x 4.6 mm and 4.6 µm of pore size). Mobile phase was composed of solvent A and B (described in the previous section). The program method was set up in gradient mode with a variation of 5% to 95% of solvent B during 30 min with a 1 mL min^-1^ flow. Detection was carried out at dual mode (wavelengths at 220 nm and 280 nm) and the oven temperature was maintained at 40°C. After obtaining the peptide chromatographic profiles and identifying the peak of interest, purification was also performed by HPLC, using a C18 reverse phase semi-preparation column (Shimadzu, dimensions of 250 × 20 mm and 15 µm of pore size). Solvents A and B were used as a mobile phase at a flow of 5 mL min^-1^. The program method was carried out in gradient mode with a variation of 30% to 60% of solvent B in 90 min, at 40°C. A peptide mass of 60 mg was dissolved in 5 mL of solvents A: B (70:30, v/v) and injected manually into the system. Detection was performed in dual mode (220 and 280 nm). The chromatogram was monitored, and the peptide fractions were collected using an automatic fraction collector (Shimadzu, model FRC-10A). Subsequently, the fractions collected were analyzed in analytical mode using the same conditions preformed for the chromatographic profile of crude peptide, obtaining the pure fractions of the Ctx(Ile^21^)-Ha peptide.

Liquid Chromatography coupled with Mass Spectroscopy (Shimadzu UFLC, Prominence/Bruker Amazon SL) were performed with the same parameters as analytical HPLC analysis, at a flow rate of 0.5 mL min^-1^, for determination of mass/charge ratio of the peptide and further confirmation of the Ctx(Ile^21^)-Ha peptide obtaining.

2.3. **Production of coted-microparticles**
     2.3.1. **Production of microparticles (E)**

The ionic gelation method was employed for microparticles production. The solutions, or dispersion phase, containing the Ctx(Ile^21^)-Ha peptide (0.2 g L^-1^ and 0.4 g L^-1^, respectively) were added in a 2% (w/w) sodium alginate solution, using an UltraTurrax-T18 (IKA- Labortechnik, Germany) at 25,000 rpm min^-1^ and ultrasonicated with an ultrasound probe (Ultrassonic Processor, Hilscher, Germany) for 30 min. Another solution, or crosslinking solution, containing 5% of aluminum chloride solution was prepared. The syringe was loaded with the dispersion solution (20 mL) and a syringe pump was used (New Era Pump System Inc., USA), at an infusion rate of 1.2 µL h^-1^. Microcapsules were produced dropwise into the crosslinking solution at RT. Two needle diameters were used (1.1 and 2 mm, capsules labeled ‘H’ and ‘C’, respectively). After that, they were dried in a drying oven at 40 °C for 4 h and stored in the dark until their characterization and application.

2.3.2. **Fluidized-bed coating of (ER) microparticles**

The fluidized bed coating method was used for coating. The microcapsules were placed in a fluidized bed (LabMaq MLF 100, Brazil) at 50°C, 0.2 L min^-1^ blower, 0.4 mL min^-1^ peristaltic pump and 70% vibration as system conditions, with 40 mL of coating solution (CS). CS was prepared with 10% HPMCAS, w/w, 25% ammonium hydroxide, w/w, 2.5% triethyl citrate, w/v, and 62.5% water, w/v. Samples were labeled for all analyzes as ‘C1’ (0.2 g L^-1^ of Ctx peptide encapsulated and coated with HPMCAS), ‘C2’ (0.4 g L^-1^ of Ctx peptide encapsulated and coated with HPMCAS), ‘H1’ (0.2 g L^-1^ of Ctx peptide encapsulated and coated with HPMCP), ‘H2’ (0.4 g L^-1^ of Ctx peptide encapsulated and coated with HPMCP), ‘BR’ (microparticle control without peptide).

2.4. **Characterization of coated-microparticles**
     2.4.1. **Encapsulation efficiency**

To estimate the encapsulation efficiency, measured by an indirect method, the total amount of free peptide in the crosslinking solution after microencapsulation per mL was measured. To corroborate the encapsulation efficiency of microencapsulation process, it was measured using a method previously described, with some modifications and adapted for this system (Di Giorgio et al., 2019; Ghatak & Iyyaswami, 2019), using the Equation 1.

$Encapsulation efficiency \left( EE \% \right)=\frac{TP-FPC}{TP} X 100$	(Equation 1)

Where *TP* is the total Ctx peptide content supplied and *FPC* is the free peptide content of microcapsules. To quantify the *EE*, 50 mg of the microcapsule were dissolved in a 5% sodium citrate solution, w/v, at 180 rpm for 3 h. Then, the peptide was rapidly extracted with solvent A and B for 2 h and centrifuged. For the analysis, 1 mL of solution was read at 280 nm in a UV spectrophotometer (Shimadzu UV1800, Japan), and calculated using as equivalent parameter the tryptophan absorption in this wavelength, using a molar absorptivity of 5,690 M^-1^ cm^-1^.

2.4.2. **FT-IR/FT-Raman analysis**

The FT-IR spectra of the coated microcapsules were analyzed using a spectrometer (Perkin-Elmer, Frontier model, USA) using the attenuated total reflectance (ATR) fixture (Bruker Vertex 70 FTIR). The transmittance spectra of the microencapsulated powders were documented with a resolution of 4 cm^-1^. Raman spectroscopy was analyzed using a Burker Ram II Raman spectrophotometer, 1,064 nm laser, 200 mW power and Ge detector. All experimental data were analyzed in the 4,000 to 400 cm^-1^ range using OriginPro 2019b software.

2.4.3. **Morphological analysis**

The morphological properties of the coated-microencapsulated peptide were carried out using a Scanning Electron Microscope (FE-SEM) (Jeol Ltd., Japan). The microcapsules were coated with charcoal after bonding with the ends of the tape. Surface and cross-sections of the samples were analyzed with 10µA, 2.0 kV energy emission current, current probe of 9, 6.5 mm working distance, Secondary Electron Imaging (SE) mode, 15,000x magnification (1 pixel = 0.935 nm).

2.4.4. **Thermal analysis**

The thermogravimetric (TGA) and differential scanning calorimetry (DSC) study was performed with the microparticles, pure peptide and microparticle without peptide, using between approximately 10 – 12 mg of sample, wrapped in an alumina crucible and stored under a synthetic air atmosphere (injection rate of 100 mL min^-1^), and evaluated from 30 to 600°C (5 °C min^-1^). The results that were obtained with DSC-TGA Equipment (SDT Q600 V20.9 Build 20, TA Instruments) and processed with the Universal Analysis 2000 software.

2.4.5. **X-Ray Diffraction analysis**

The crystallinity of the microcapsules was studied using an X-ray diffractometer (XRD-6000, Shimadzu), at 40 kV and 30 mA, using CuKα1 and CuKα2 radiation sources (λ = 1.5406 and 1.5444 Å, respectively). Continuous scanning in the range of 2θ from 10° to 80° at 2° min^-1^, using an aluminum or glass sample holder depending on the amount of sample available.

2.5. ***In vitro* release profile**

The SG solution was prepared with a volume of 10 mL of HCl 0.1 mol L^-1^ at pH 3.5, adjusted with HCl 1 mol L^-1^. Then, 1 mL of SG was collected after 1, 2, 5, 10, 15 and 20 min and measured at a UV spectrophotometer at a wavelength of 280 nm; ii) s*imulated intestinal fluid (SIF)*, procedure that mimics the duodenum and jejunum (first and second portions of the small intestine of poultry). Microcapsules samples were weighted and a mass of 46.75 ± 4.92 mg of them were placed in an SI solution at 42°C for 60 min. The SI solution was prepared with 10 mL of HCl 0.1 mol L^-1^ at pH 6, adjusted with NaOH 1 mol L^-1^. Then, 1 mL of SI was collected after 1, 2, 5, 10, 15, 20, 30, 45 and 60 min and measured at a UV spectrophotometer at a wavelength of 280 nm.

2.6. **Hemolytic activity assay**

For this test, erythrocytes separated from plasma by centrifugation (10 min at 1,600 rpm) were used and washed with PBS buffer pH 7.4. Then, a 2% (v/v) cell suspension of human erythrocytes was incubated in PBS buffer with each of the 4.29 ± 0.27 mg mL^-1^ microparticle formulations at 37°C with stirring (100 rpm). Aliquots were taken at different incubation times (1, 4, 8 and 24 h) and centrifuged (10 min at 2,300x g). As a positive control (hemolytic activity = 100%), a solution of 1% (v/v) of Triton X-100 detergent was used. Hemolysis was analyzed from the absorbance values of the supernatant at 570 nm, according to the Equation 2:

$Hemolysis \left( \% \right)=\frac{\left( A_{s} - A_{b} \right)}{(A_{t} - A_{b})} x 100$	(Equation 2)

Where *A_s_* is the absorbance of the samples, *A_b_* is the mean of the absorbance in the buffer and *A_t_* corresponds to the total lysis of the cells. All procedures were performed in triplicate.

3. **RESULTS**
   3.1. **Ctx(Ile^21^)-Ha peptide analysis**

For a reliable peptide analysis, it is necessary to use the molecule with purity higher than 95% or it must be purified (Ferreira Cespedes et al., 2012). The antimicrobial peptide Ctx(Ile^21^)-Ha was successfully obtained, with more than 95% purity. The characteristic peak of the peptide was obtained by analytical HPLC with a retention time of 18.82 min (Figure 1A). In the spectrum, the peaks of 573 (Z = +4), 763.6 (Z = +3) and 1,144.9 (Z = +2) represent the peaks of the mass/charge ratio of a molecule with a molecular weight of 2,289.72 g mol^-1^, which is exactly the theoretical molecular weight of the antimicrobial peptide Ctx(Ile^21^)-Ha, confirming its successful obtaining and characterization (Fig. 1).
